# Response to CAR T cell therapy can be explained by ecological cell dynamics and stochastic extinction events

**DOI:** 10.1101/717074

**Authors:** Gregory J. Kimmel, Frederick L. Locke, Philipp M. Altrock

**Author notes:** These authors contributed equally (Frederick L. Locke, Philipp M. Altrock). Corresponding authors, H. Lee Moffitt Cancer Center, and Research Institute 12902 USF Magnolia Drive, Tampa, FL 33612, USA.

## Abstract

Chimeric antigen receptor (CAR) T cell therapy is a remarkably effective immunotherapy that relies on *in vivo* expansion of engineered CAR T cells, after lymphodepletion by chemotherapy. The laws underlying this expansion and subsequent tumor eradication remain unknown. Still, about 60% of CAR T-treated patients are likely to progress; their tumors are not eradicated. Here we seek to understand and disentangle the multiple processes that contribute to CAR expansion and tumor eradication. We developed a mathematical model of T cell-tumor cell interactions, and demonstrate that CAR T cell expansion is shaped by immune reconstitution dynamics after lymphodepletion and predator prey-like dynamics. Our cell population model was parameterized using patient population-level data over time and recapitulates progression free survival. As an intrinsic property, we find that tumor eradication is a stochastic event. Our cell population-based approach renders CAR T cell therapy as an ecological dynamic process that drives tumors toward an extinction vortex. Even if a clinical event, such as progression, is likely, its timing can be highly variable. We predict how clinical interventions that increase CAR T memory populations could improve the likelihood of tumor eradication and improve progression free survival. Our model can be leveraged to propose new CAR composition and dosing strategies, assess the need for multiple doses, and identify patient populations most likely to benefit from CAR T with or without additional interventions.

## INTRODUCTION

Relapsed and refractory large B cell lymphoma (LBCL) is the most common subtype of Non-Hodgkin Lymphoma, the most common hematologic malignancy in the US with 72,000 new cases (4.3% of all cancer) and 20,000 deaths (3.4% of all cancer deaths) in 2017^1^. In LBCL patients that do not respond to chemotherapy, the median overall survival is under 7 months^2^. These patients could benefit from autologous chimeric antigen receptor (CAR) T cell therapy that uses genetically engineered T cells re-targeted to CD19, a protein specific to B lineage cells from which LBCL arises^3^. ZUMA-1 was a pivotal, multi-center, phase 1-2 trial of axicabtagene ciloleucel (axi-cel), a second generation CD19-directed CAR T cell therapy with CD28 co-stimulation, for chemo-refractory LCBL patients (n=101 patients treated)^4–6^. Overall Response Rate (ORR) and Complete Response (CR) rate were 82% and 54%—respective responses would have been 26% and 7% with standard chemotherapy^2^. The US FDA approved axi-cel for relapsed/refractory LBCL as treatment for adults in 2017. Similar results were demonstrated with other CD19 CAR T cell therapies^7,8^. Patients not achieving CR or partial response (PR) by 90 days after the start of CAR T treatment fare poorly. Overall, 60% of LBCL patients eventually progress although many of them see a temporary reduction in tumor burden.

Quantitative models of CAR T dynamics and tumor responses are needed to better predict patient outcomes, and to approach the modeling of combinatorial therapies. The utility of mathematical modeling of anti-cancer combination therapies has been established for untargeted, targeted and combination therapies^9–12^—applying eco-evolutionary^13,14^ and cell-differentiation^15,16^ theories. Cellular immunotherapies, such as CAR T cell therapy, describe a new frontier for predictive mathematical modeling of treatment outcomes^17^, largely because the cellular drug itself does not follow conventional dynamical patterns. Recent works used mixed-effect modeling of CAR-T cell therapy^18^ to model the expansion of the CAR T cell drug tisagenlecleucel^19^, in combination with therapies that treat cytokine release syndrome. Interestingly, this model assumed a CAR T cell compartmentalization as an explanatory approach to the complex CAR dynamics over time. Other recent works considered eco-evolutionary dynamics to explain CAR T cell expansion and exhaustion^20^, and signaling-induced cell state variability^21^, both inferred from *in vitro* data and without considering interactions between T cells and tumor. Here, we seek a fundamental understanding of T and CAR T cells including tumor cell dynamics *in vivo*.

Tumor response and survival, and toxicities (and cytokine release syndrome) to a lesser extent^22^, have been linked to a maximum in CAR T cell density over time^4^. Furthermore, *in vivo* CAR T cell expansion, tumor response, and toxicity are impacted by CAR T differentiation states^23^, T cell-competition dynamics^24^, and homeostatic signals^25^. In addition, T cells secrete signals that regulate their proliferation in a density-dependent way^26^, and stimulate or inhibit differentiation^27^. Finally, the amount of cognate antigen encountered, especially the CD19 positive tumor, plays a dynamic role in T cell turnover because it directly influences T cell differentiation into effector cells, and thus indirectly regulates CAR expansion and life span^28^.

We analyze clinical data of T and CAR T expansion and tumor response. We focus on two CAR T cell differentiation states of naïve/long-term memory and effector cells^29,30^. We propose that co-evolution in the T cell homeostatic niche drives the observed CAR T cell dynamics and imply tumor/antigen feedback onto CAR T cell differentiation into effector cells^31^. T cell proliferation, differentiation (into effector cells), and CAR T cell survival are key processes that bear characteristics of eco-evolutionary dynamics. An improved understanding of the ecological interactions and dynamics involving lymphocytes and tumor cells within patients during CAR T cell therapy could be leveraged to explore new dosing and combination therapies. We integrate and model *in vivo* normal T and CAR T cell dynamics (memory to effector cells) together with changes in patients’ tumor burden, in order to discover new principles that drive durable response or relapse.

## METHODS

We developed a mathematical modeling framework that describes dynamics and interactions among normal T cells, CAR T cells, and tumor cells. The model considers four cell populations in the form of continuous-time birth and death stochastic processes and their deterministic mean-field equations: normal naïve/memory T cells, *N*, naïve/memory CAR T cells, *M*, effector CAR T cells, *E*, and antigen-presenting tumor cells, *B*. We assume that memory cells grow toward a homeostatic maximum, reflected by the carrying capacities *K_N_ and K_M_*.

We consider the case of lymphodepleting chemotherapy prior to infusion of autologous CAR T cells. We set time to 0 at the time of CAR infusion. Normal and CAR memory T cell populations then grow toward their respective carrying capacities and influence each other. This mutual influence gives rise to co-evolution. Normal memory cells expand at the rate *r_N_\**log[*K_N_*/(N+M)]. CAR memory cells, for our purposes labeled as CCR7+^29,31,32^, expand at the rate *r_M_\**log[*K_M_*/(N+M)]. Of note, we allow for a time-dependent intrinsic growth rate of memory CAR T cells around time τ > 0, 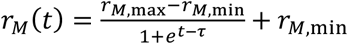. This transition from a large to a smaller growth rate *r_M_* allows for exogenous control of the competitive advantage/disadvantage of CAR T cells, potentially as the effects of lymphodepletion decay.

Memory CAR T cells differentiate into effector CAR T cells, and become CCR7-^29,31,32^, at a rate *r_E_*(B) and depending on the amount of antigen CD19. In the model, we exclude possible influences by other sources of CD19, such as normal B cells. In ZUMA-1, ~50% of patients did not have detectable normal B cells in circulation at the time of lymphodepletion, and 3 months after CAR T infusion under 20% of patients had detectable normal B cells. Thus, in the patient population we are modeling, normal sources of CD19 do not seem to play a large role during the time CAR T cells are most active, and we used the following antigen-driven, piecewise-linear feedback of tumor mass on the rate at which effector CAR T cells emerge: *r*_*E*_(*B*) = *r*_*E*_(0)(1 + α_1_ min{*B*/*B*_0_, α_2_}).

While we assume that memory CAR T cells could persist indefinitely (possibly at very low concentrations), effector CAR T cells have a maximum life-span determined by their intrinsic death rate *d_E_* and by the fact that interaction with CD19+ tumor cells also leads to exhaustion and subsequent effector cell death at rate *γ*^33^.

The tumor cell population *B* grows autonomously at the net growth rate *r_B_*, and experiences tumor killing at rate *γ_B_*, proportional to the number of CAR T effector cells *E*. This dynamical system can be written, in the mean-field limit, as a set of deterministic ordinary differential equations:

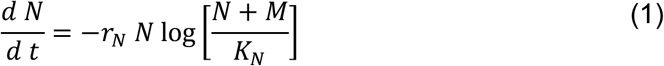

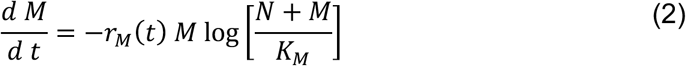

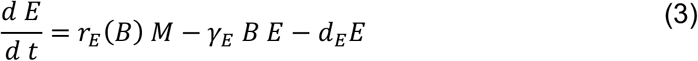

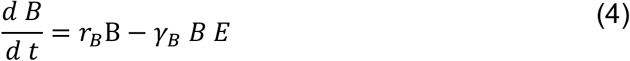

For a derivation of these equations and their associated stochastic birth and death process, see Supplementary Information (SI), where we present a stability analysis. Fig. 1 A shows a schematic of this complex dynamical system, Fig. 1 B summarizes the possible events at the cellular level. The qualitative, clinically important dynamics, ranging from no response, to transient response followed by relapse, to long-term response, are shown in Fig. 1 C-E.

**Figure 1:**
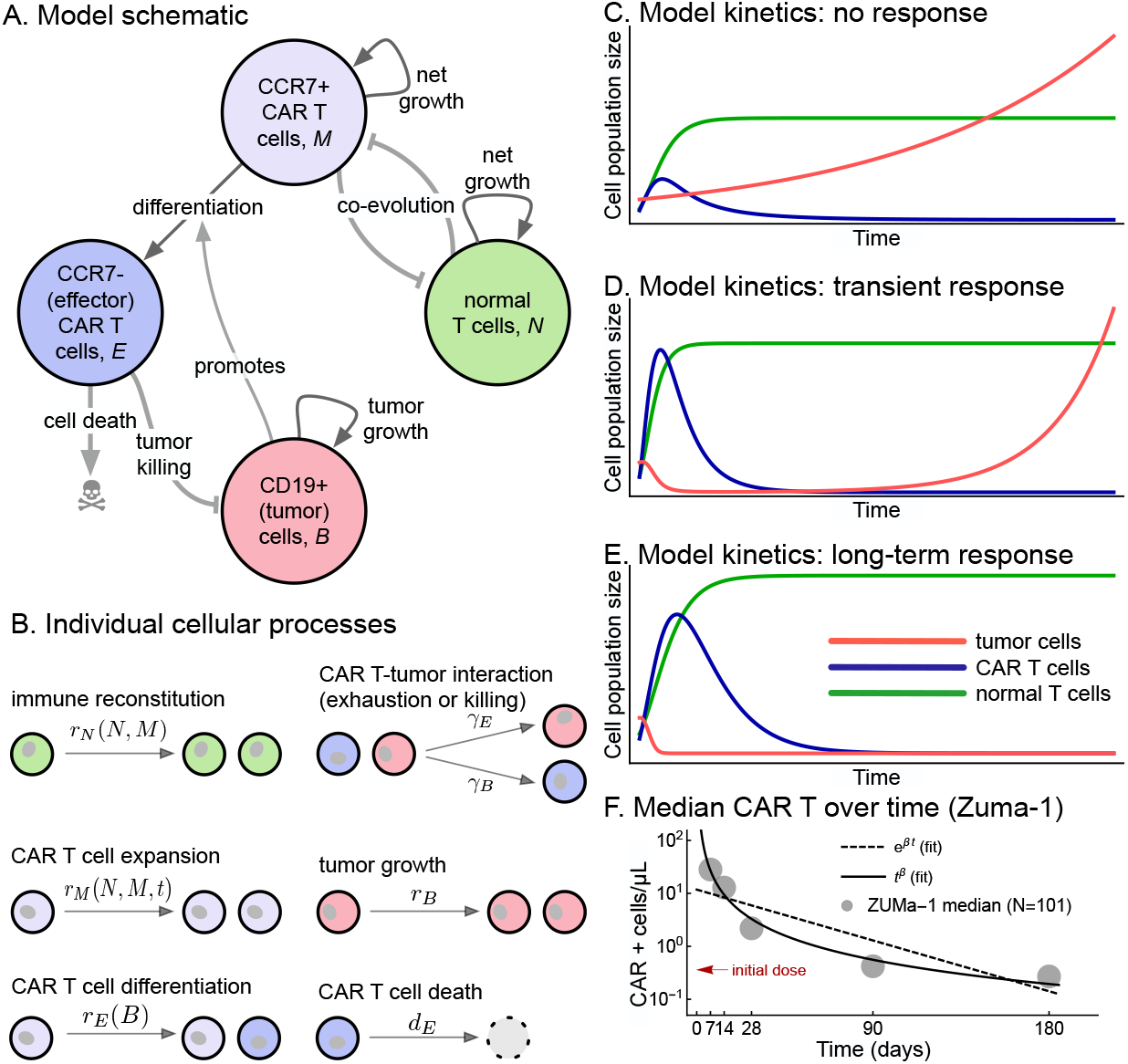
T cell interactions, CAR T cell compartmentalization, and tumor feedback on CAR T cell differentiation. **A**: Model schematic, assuming four cell compartments: memory (CCR7+) CAR T cells, M, proliferate and engage with resident lymphocytes, N (depleted by lymphodepleting chemotherapy), and differentiate into effector CAR T cells (CCR7-), E. E cells of finite life span engage in killing CD19+ tumor and other B cells. Their production is impacted by CD19. **B**: On the level of individual cells, this system results in six cellular kinetic reactions. We seek to explain the important patient kinetics of no response to CAR T cell therapy (**C**), transient response of initial tumor decline followed by progression/relapse (**D**), and long term/complete response (tumor is eradicated) (**E**). These example dynamics were generated using Equations (1)-(4) with hand-picked parameters. **F**: Median CAR T positive cells per μL peripheral blood (symbols, from ZUMA-1^6^), vs. regression fits according to exponential or power-law decay (details see SI).

### Clinical data integration

We used population-level data from LBCL patients treated on the multi-center ZUMA-1 trial^6^, summarized in Fig. 2 A. The ZUMA-1 trial reported quartile CAR T cell levels in peripheral blood over five consecutive time points (days 7, 14, 28, 90, 180), which were used to parameterize the mean-field model on the median population level (see SI). We used normal complete blood count-derived absolute lymphocyte counts (ALC, approximating the total T cell compartment), measured in patients receiving CAR T cell therapy (days 0, 5, 7, 14, 28, 90, 180). These counts include CAR T cell density, yet differences in parameters estimates were minimal and no changes in homeostatic ALC were detected, see Fig. 2 B.

**Figure 2:**
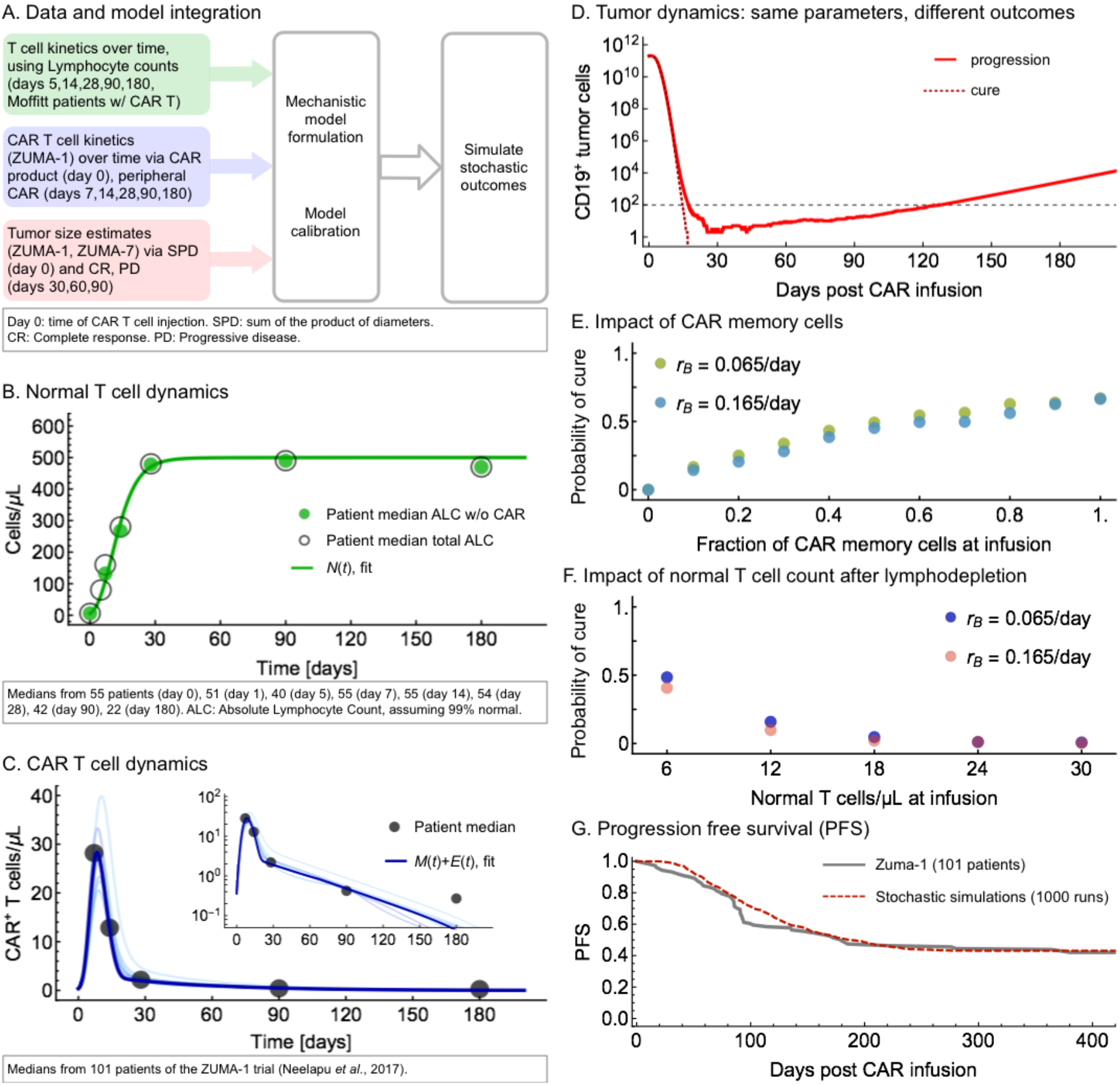
A population-level model of T cell co-evolution, complex CAR T cell dynamics can predict clinical endpoints as stochastic events. **A**: Schematic of population-level data integration to parametrize the mathematical model presented in Fig. 1; we used longitudinal data of peripheral absolute lymphocyte count (ALC), peripheral CAR positive cell counts per µL, and the tumor size changes as estimated from patients of the ZUMA-1 trial with complete response (CR) or progressive disease (PD). We assumed that, at days 30, 60, or 90, CRs had no detectable tumor mass, and that PDs had twice their initial tumor mass, Median initial tumor mass was 200 cm^3^. **B**: ALC was used to estimate the dynamics of Eq. (1), which did not change dramatically based on ALC or ALC subtracted CAR data. **C**: CAR positive T cell dynamics (Eqs. (2), (3)) were estimated using ZUMA-1 trial data of peripheral CAR counts and can explain peak and decay of CAR. **D**: Two example trajectories of tumor burden over time, using identical parameters and a stochastic process to model tumor extinction. Both examples enter the stochastic region (<100 tumor cells), but one escapes this extinction vortex, leading to progression. **E**: Increasing the fraction of initial memory CAR T cells (CCR7+) could improve chances of cure. **F**: Initial ALC of six cells/μL at CAR administration due to lymphodepletion is crucial; increasing this number would monotonously decrease the chances of cure. **G**: Progression free survival (PFS) was recorded in ZUMA-1 (gray line). Our stochastic model recapitulates this curve using stochastic simulations with a mixture of parameters drawn from a normal distribution with a variance of 10% of the mean (Methods, SI), and recording progression when 2 times the initial tumor mass is reached. All parameter values used are given in **Table 1.** All probabilities estimated used 1000 stochastic simulations with the same initial conditions.

All peripheral measurements of cells/μL were converted to total T cell counts assuming, 5L of blood and multiplying by 10^8^, under the assumption that 1% of cells are in periphery at all times and converting μL to L. The normal T cell carrying capacity, *K_N_* = 500 cells/µL in patients, was estimated from data by Turtle *et al*.^34^

Median tumor volume at day 0 was determined by assuming a spherical lesion, which corresponds to a volume of *B_0_*=200cm^3^. This was then converted to 2.0*10^11^ cells as per our own estimates from patients treated at Moffitt within ZUMA-1, assuming that 1 cm^3^ contains 10^9^ cells^35^. Complete response (CR) was counted as tumor size *B(t)*=0. Stable disease (SD) was converted to the tumor returning to initial size, *B(t)*=*B*_0_. Progressive disease (PD) was counted as twice the initial tumor size, *B(t)*=2\**B*_0_. Patients in none of these categories after 1000 days would be defined as undetermined, however none of our simulations led to undetermined patients.

### Parameter estimation

We obtained *median* parameter values using a nonlinear least squares regression function with the data-variance as weights. The nonlinear optimization problem used for data fitting was solved using the BFGS algorithm from the optimization package in Julia, wrapped around solving the 4D differential equations^36,37^. Data that was only available in quartiles: we assumed an approximately normal distribution and estimated the variance from the difference between the quantiles. We generated fits to the available quartiles of CAR T cell densities measured at the five consecutive time points available. We fit parameters in equations (1)-(2) first, as these decouple from the other equations. Then we fit effector CAR T cell and tumor cell dynamics, equations (3)-(4) (for details, see SI). The parameter values obtained are shown in Table 1, and their good agreement with median clinical data is shown in Fig. 2 B for the normal T cell dynamics, and in Fig. 2 C for the CAR T cell dynamics.

**Table 1:** Parameter and initial condition values as identified by our machine learning procedure and literature search, also see Supplementary Information (SI).

| *Biological parameter* | *Symbol* | *Fitted value* | *Stochastic simulation* | *Reference* |
| --- | --- | --- | --- | --- |
| Normal T cell carrying capacity | $K_N$ | $5.00 \times 10^2$ cells/μL | $2.50 \times 10^{11}$ cells | <sup>34</sup> |
| CAR memory T cell carrying capacity | $K_M$ | $1.60 \times 10^2$ cells/μL | $8.00 \times 10^{10}$ cells | this work |
| Normal T cell net growth rate | $r_N$ | $1.60 \times 10^{-1}$ day<sup>-1</sup> | unchanged | this work |
| Initial CAR memory net growth rate | $r_{M,max}$ | $4.14 \times 10^{-1}$ day<sup>-1</sup> | unchanged | this work |
| Final CAR memory net growth rate | $r_{M,min}$ | $2.15 \times 10^{-2}$ day<sup>-1</sup> | unchanged | this work |
| CAR memory net growth rate switch | $\tau$ | $1.88 \times 10^1$ days | unchanged | this work |
| Basal CAR asymmetric diff. rate | $r_E(0)$ | $2.26 \times 10^0$ day<sup>-1</sup> | unchanged | this work |
| Effector CAR death rate | $d_E$ | $2.93 \times 10^{-1}$ day<sup>-1</sup> | unchanged | this work |
| Tumor net growth rate | $r_B$ | $6.50 \times 10^{-2}$ day<sup>-1</sup> | unchanged | <sup>57</sup> |
| Effector CAR exhaustion rate | $\gamma_E$ | $3.00 \times 10^{-3}$ day<sup>-1</sup>cm<sup>-3</sup> | $3.00 \times 10^{-12}$ (days*cells)<sup>-1</sup> | this work |
| Tumor killing rate (by effector CAR) | $\gamma_B$ | $3.00 \times 10^{-2}$ μL day<sup>-1</sup>cell<sup>-1</sup> | $6.00 \times 10^{-11}$ (days*cells)<sup>-1</sup> | this work |
| Increased asymm. diff. rate | $\alpha_1$ | $4.29 \times 10^0$ | unchanged | this work |
| Max tumor ratio | $\alpha_2$ | $1.35 \times 10^0$ | unchanged | this work |
| Initial median normal T cell number | $N(t = 0)$ | 6.00 cells/μL | $3.00 \times 10^9$ cells | <sup>5</sup> |
| Initial CAR T cell number | $M(0) + E(0)$ | 0.36 cells/μL | $1.80 \times 10^9$ cells | <sup>5</sup> |
| Initial median tumor cell number | $B(0)$ | 200 cm<sup>3</sup> | $2.00 \times 10^{11}$ cells | <sup>5</sup> |

### Model analysis and stochastic time-forward simulations

We analyzed the deterministic mean field model using linear stability analysis (see SI). In the mean-field limit our parameters lead to eventual cancer progression in every patient. However, many trajectories spend large amounts of time *near* the tumor-free (*B* = 0) state.

Small populations cannot be adequately captured with a deterministic model (see SI), but that populations can be driven toward an extinction vortex—near small population sizes, stochastic dynamics gain increased importance^38^. This is exemplified in Fig. 2 D, where the exact same set of parameters lead to either tumor extinction or progression.

Fully stochastic simulations with a system comprised of billions of cells would take days or weeks to run for a single patient. This is computationally prohibitive since many patient trajectories are needed to gather statistics. To alleviate this issue while maintaining the accuracy of the stochastic model, we developed a hybrid model that explois the fact that only certain populations become small and that their fluctuations should only be relevant when a population is below a given threshold. The hybrid model operates in the deterministic limit if the tumor population is above a threshold of 100 cells and updates stochastically if below the threshold. The other populations are updated deterministically in parallel. Since we are only interested in whether the tumor is eradicated, our model only switches the tumor between deterministic and stochastic model, all other populations remain deterministic.

We ran the deterministic system using the built-in solver Rosenbrock23 of the *DifferentialEquation* package in Julia, and simulated the master equation using a Gillespie algorithm^39^. For a typical simulation result with stochastic outcome, we simulated 1000-10000 virtual patients, each drawn with their own, slightly different mixture of parameters. Parameter variations were achieved using a normal distribution with mean chosen as the median parameter value estimated from model fitting (described above), and variance taken as a fraction of that mean (typically between 5-15%). A patient was counted as cured when their tumor reached 0 cells, which is an absorbing state of the stochastic birth and death process. A patient was counted as progressed when their tumor mass hit the threshold of 5 times their mass prior to treatment. This high threshold was used to avoid the occasional patient’s tumor that grows rapidly enough to reach a smaller threshold before decreasing due to treatment. Observed differences in progression times due to alternative tumor size used for progression, e.g. 1.2, 2.0, or 5.0 times the initial tumor burden, are relatively small (about 6%), the final outcome (cure or progression) was unchanged. These two endpoints led to Kaplan-Meier progression free survival (PFS) curves for simulated patient, generated by noting the time of progression.

## RESULTS

### Complex dynamical processes shape CAR T cell expansion and persistence

In Fig. 1 F, we show that no simple exponential decay can explain CAR T cell dynamics *after* peak. Rather, a power law of the form *t^−β^* (as opposed to e^−^*^βt^*) with a suitable parameter *β* can statistically describe CAR decay, which indicates that multiple time scales *within* the CAR T cell population are relevant. Furthermore, CAR expansion up to the peak is important. A decay process alone cannot describe expansion, pointing to the impact of additional complexity. Here, this additional impact was assumed to stem from interactions among normal and CAR T cells during return to homeostasis, which we integrate using an ecological co-evolutionary framework.

### Co-evolution among normal T and CAR T cells and memory to effector CAR T differentiation explain clinical outcomes

All patients received lymphodepleting chemotherapy followed by autologous CAR T cell infusion. The degree of CAR T cell expansion has been associated with durable response to therapy^4,18^. Two key components of our model are mechanisms that drive CAR T expansion and duration of persistence: (a) external homeostatic signals (normal T-CAR T interactions), (b) CAR memory to effector differentiation rate, which is proportional to tumor mass. This model can qualitatively describe the three different clinical trajectories of no response, transient response and long-term response (Fig. 1).

The dynamical system is hierarchical in the CAR T cell compartment. Memory T cell compartments influence each other and impact more differentiated compartments, but not vice versa. This means that one can establish quantitative rules of normal and CAR T cell interactions, independent of effector CAR T cell and tumor properties. We originally fit a generalized logistic model and found that the optimization routine selected a Gompertzian growth model in the non-tumor killing T cell compartments^40^. A possible mechanism for the emergence of Gompertzian growth was recently discussed in cancer population growth emerging from an interaction-saturation process as the population increases in size and diversity^41^. In an analogous fashion, as the T cell population returns to homeostatic levels, antigen-experienced phenotypes are filled, leading to proportional slowing of growth during immune reconstitution. The resulting decrease in available cell states can explain a characteristic logarithmic approach of the T cells’ carrying capacity.

The CAR T system quickly returns to near-homeostatic values, with 32% normal T cells at day 7, and 94-98% after day 28 (Fig. 2 B, assuming normal density of about 500 cells/μL blood^24^). We found that the CAR T cell carrying capacity is lower than its normal T cell counterpart.

Memory CAR T cell dynamics are best explained using a transition from a rapid to a slow growth phase around day 19 post CAR injection, and memory CAR to effector CAR T cell differentiation is positively, but weakly, affected by tumor mass. We estimated that the CAR T cell carrying capacity is reduced to about 33%. Due to rapid expansion into effector CAR T cells, and an initially higher CAR T cell proliferation rate, the overall CAR+ cell populations are non-monotonic and exhibit a peak within the first two weeks post-infusion. We thus understand the process as follows. In the short term, memory CAR cells and effector cells expand, but they are replaced by normal T cells in the long term.

### Tumor eradication by CAR T cells is deterministically unstable

The four-dimensional deterministic system was analyzed to determine its steady states and their stability. Two long-term outcomes are important (see SI discussion). First, normal T cells return to homeostatic levels, while CAR T cells vanish, and the tumor eventually grows. This state was stable if the normal T cells’ carrying capacity was larger than the CAR T cells’ carrying capacity.

Second, the tumor was brought to a stable value or it was eradicated. However, the two corresponding steady states were stable only if CAR T cells’ carrying capacity was larger than that of normal cells, which is highly unlikely. Hence, in clinical settings where CAR T cells are slightly or strongly maladapted in the long run, tumor eradication is deterministically unstable. Nonetheless, the non-monotonic CAR kinetics indicates that the tumor mass often shrinks at least for some time.

### Stochastic tumor extinction can be an explanation for observed treatment success rates

As a result of the fact that tumor extinction is not possible in the deterministic large population size regime, we propose that malignant B cell extinction is a stochastic event. Stochastic extinction results from the tumor being driven close to an extinction vortex^42^. This means that, even if cure is likely under clinically favorable conditions (and progression unlikely), the time at which cure (i.e. eradication of tumor) becomes certain follows a potentially broad distribution.

The probability of tumor extinction can be calculated as a function of specific model parameters for a fixed point in time, or for all times (Fig. 2 E, F). Treatment success critically depends on a sufficient fraction of long-term memory/naïve (CCR7+) cells in the CAR product (Fig. 2 E), and on the effectiveness of lymphodepletion that reduce absolute lymphocyte counts (Fig. 2 F).

For a fixed set of parameters (Table 1), tumor eradication can be tracked over time, as shown in Fig. 2 G. Remarkably, although we had not used progression free survival directly as a goal function to find suitable model parameters (see SI), our calibrated stochastic model can recapitulate progression free survival (PFS) of the ZUMA-1 trial.

### Stochastic simulations can be used to estimate the influence of the parameters on PFS

The stochastic representation of the ecological CAR T cell dynamics allows us to calculate survival curves from a ‘virtual cohort', with each patient defined by a unique combination of randomly chosen parameter values. We focused on the two scenarios of transient and long-term response (sketched in Figs. 1 D, E).

The ‘typical’ case in our simulations, defined by using all median parameter values (Table 1), does not progress until day 180 post CAR infusion. We were thus interested in the overall impact of parameter variation on survival outcomes (Fig 3 A), assuming that all parameters are normally distributed around the mean value for the median patient, with a variance calculated as a small fraction of the mean. Increasing parameter variance led to more heterogeneous PFS curves. All differences between simulated PFS curves were statistically significant using a log-rank test due to the large simulated cohort size; hence, we focus on the magnitude of change of PFS, e.g. at a specific point in time. Our PFS curves show an initial plateau, which stems from the fact that at least some part of the tumor responds to CAR T cell treatment initially. In the longer term, the PFS curves reflect the inherent stochasticity of tumor extinction or escape.

**Figure 3:**
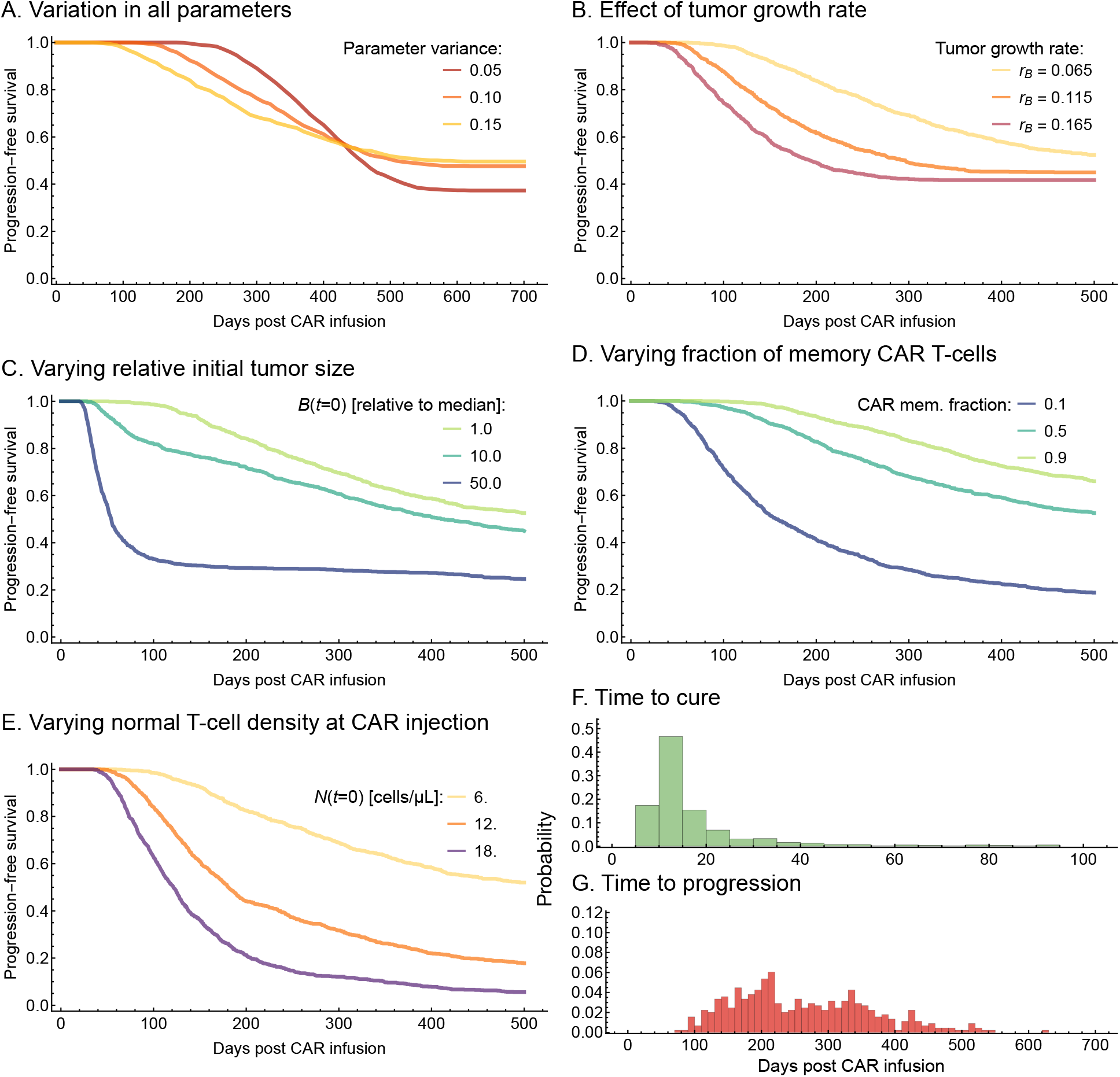
The mathematical model recapitulates and predicts progression free survival (PFS), and can suggest actionable therapy improvements. 1000 simulated patients were used to generate each PFS curve. **A**: Impact of parameter variation on the PFS. All subsequent panels use σ = 0.15. **B**: Impact of tumor growth rate on the PFS. **C**: Impact of initial tumor size on the PFS. Much larger tumors lead to some patients progressing earlier since the CAR could not expand fast enough. **D**: Impact of CAR T infusion phenotype composition. In general, a higher memory fraction lead to better PFS rates. **E**: Impact of lymphodepletion on PFS. Similar to Figure 2G, a sizable impact on PFS is observed by doubling or tripling the amount of normal T cells after depletion. **F**: The distribution of cure times for the median parameters. Most patients are cured before day 100. **G**: The distribution of progression times for the median parameters. Most patients progress between days 80-400. All parameter values used are given in **Table 1.** All probability estimated used 1000 stochastic simulations with the same initial conditions.

### Larger and faster growing tumors have worse PFS, with very large tumors leading to increased likelihood of rapid progression

Intrinsic tumor properties are the tumor growth rate, *rB*, and initial tumor burden (*B*_0_, where we report values relative to the median tumor size of 200 cm^3^). Tumor growth rate increases had a strong detrimental effect on PFS at early timepoints (Fig. 3 B). At day 100, PFS of slow-growing tumors (*r_B_*=0.075/day) could be above 90%, whereas PFS of fast-growing tumors (*r_B_*=0.175/day) were around 70%. In comparison to the median tumor size, ten-fold larger tumors (*B*_0_=10) showed a drop 70% in PFS at day 100 (from 90% PFS), and to 55% for fifty-fold larger tumors (*B*_0_=50, Fig. 3 C).

### PFS declines with low levels of CAR memory content and insufficient lymphodepletion

Next, we asked how PFS is influenced by properties of the CAR and normal T cell population prior to injection. To this end, we considered changes in the fraction of CAR memory fraction at day 0, and variation in the efficacy of lymphodepleting chemotherapy. In concordance with Fig. 2 E, larger differences in PFS were detected when the memory CAR T cell fraction was low (Fig. 3 D). Comparably, the potential failure to effectively lymphodeplete prior to CAR treatment could have drastic effects. Doubling the initial normal T cell population size could halve PFS at day 100 and thereafter (Fig. 3 E).

### Cure events according to direct CAR T cell predation occur early, while progression events can occur late

Our stochastic modeling revealed that cure occurs early; most simulations resulted in cure between days 10 and 35, and we rarely found late cure events up to day 100 (Fig. 3 F). Meanwhile, progression times were distributed over a broad range of time points. Typical progression times occurring anywhere between days 20 and 500 (Fig. 3 G). These large differences in time scales occur because cure, as a stochastic tumor extinction event, is much more likely to occur before the effector CAR T cell begins to decline after day 14 (compare to Fig. 2 C). Differences in timing of events also become clear by looking at more fine-grained comparison of survival at distinct time points, as a function of initial tumor burden (Fig. 4 A) and intrinsic tumor growth rate (Fig. 4 B).

**Figure 4:**
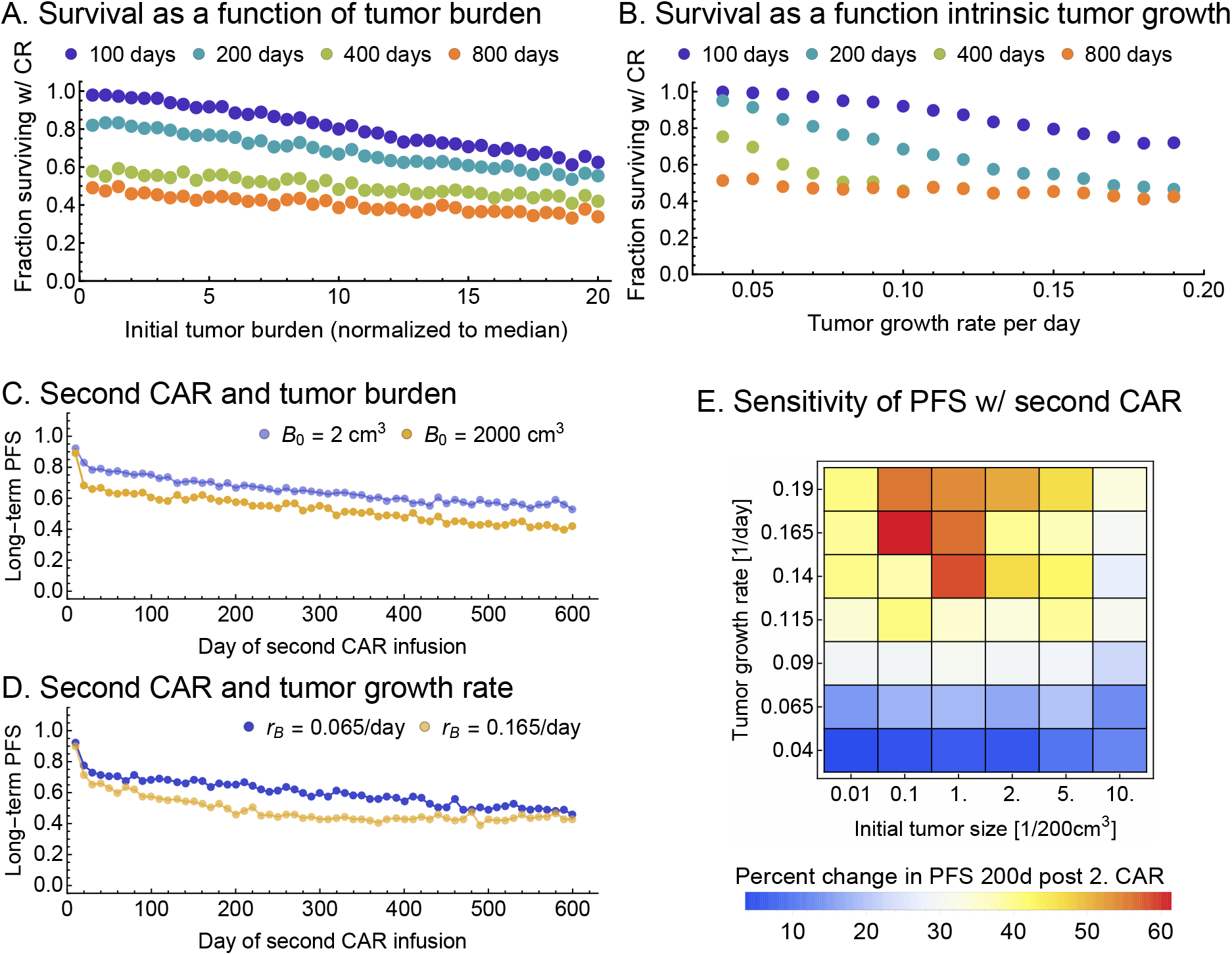
The impact of tumor size and growth rate on a second dose of CAR T cells applied after 100 days of the initial dose in regards to long-term PFS (defined at 700 days). PFS=Progression free survival. **A-B**: Fine-grained comparison of survival at distinct time points, as a function of (**A**) initial tumor burden and (**B**) tumor growth rate. **C**: The influence of initial tumor burden on the effect of a second infusion of CAR. **D**: The influence of tumor growth rate on the effect of a second infusion of CAR. **E**: The sensitivity of the PFS to initial tumor size and growth rate 200 days after initial dose and 100 days after the second dose, which was given at day 100.

### Patient population-level modeling can be used to explore the expected effects of alternative treatment strategies

Our mathematical model can be used to explore alternative treatment scenarios and strategies that were not part of previous clinical studies. For example, a second CAR T cell treatment at a specified point in time can be explored. We assumed that a second dose, equal to the first dose in total CAR T cell amount and composition was given together with an appropriately timed second lymphodepleting chemotherapy. To assess possible improvement of long-term PFS, we simulated to 2 years post second CAR injection. We then explored this statistical outcome variable as a function of the timing of the second treatment relative to the first injection. We focused on variation in initial tumor size, *B_0_*, and intrinsic tumor growth rate, *r_B_*. The number of CAR T cell infused was chosen to match the ZUMA-1 median (Table 1), and we simulated 1000 trajectories. With baseline (single) treatment of long-term durable response rate of 45% (day 200), out of 1000, 550 would progress eventually. Our single dose-results (Fig. 3 G) suggested that less than 1% of these progress after day 600, which renders the 2 year mark a useful endpoint for comparison.

We expected that the additional benefit of a second dose would decrease monotonically with its timing and approach a minimal added benefit asymptotically, similar to a second ‘independent’ treatment. Indeed, the highest long-term benefit was achieved if the second CAR were given immediately after the first (Fig. 4 C, D)—a case that might be clinically unfeasible. We asked whether there is a trade-off between later second dose and associated loss of long-term benefit. Our simulations highlight the impact of the tumor growth rate, an important between-patient variability, on progression free survival. Faster growing tumors with low single dose cure rate could benefit from second dose as early as possible. These results highlight the utility of our stochastic ecological model in the context of establishing rational for novel treatment strategies.

### What are the effects of initial tumor burden and growth rate on long term PFS giving a second CAR?

The added benefit of a second dose as a function of both tumor size and tumor growth rate is non-monotonic. For intermediate and higher tumor growth rates, there can be a tumor size or a tumor growth rate that yields maximal benefit of a second dose (Fig. 4 E). At higher intrinsic tumor growth rate, *r_B_*, the expected change in PFS varies more dramatically as a function of initial tumor burden, see Fig. 4 E, where variation in day 200 (post CAR) PFS improvement is low for *r_B_<*0.1/day, and high for *r_B_=*0.14/day and above.

## DISCUSSION

In our system, five processes are potential drivers of patient outcomes: the effects of lymphodepleting chemotherapy, the composition of the CAR T cell product (e.g. CCR7+ density), CAR T cell expansion (peak), CAR durability, and tumor burden. To better understand these processes, we developed a cell population-ecological framework that describes co-evolution between normal T, CAR T (memory and effector), and tumor cells. At high densities, a deterministic mean-field approach can be used to describe the dynamics. We established that cure, as a result of tumor extinction, cannot be captured by that deterministic model and requires a stochastic individual-based description. This approach then led to predictions and sensitivity analysis of probability of cure and PFS.

Model calibration and predictions were made with respect to data that represent median patient trajectories. At this point, we do not make patient specific personalized predictions, for which a more comprehensive data set and modeling integration approach would be needed^43^, matching multiple longitudinal data from the same patient. Instead, we give proof-of-principle that the integration of longitudinal lymphocyte counts with CAR T cell counts and changes in tumor burden can explain CAR T cell population dynamics and therapy outcomes. Our approach highlights the utility of mechanistic approaches to *in vivo* CAR T cell therapy dynamics.

Our model confirms the hypothesis that sufficient lymphodepletion is an important factor in determining durable response. We predict that increases in memory CAR T cell fractions (e.g. determined by the fraction of CCR7+ cells) result in expected survival gains. This points to other necessary dynamical quantities that affect CAR expansion and tumor killing, likely impacted by upregulation of inflammatory cytokines, such as IL-7 or IL-15^25^. Thus, future modeling that follows from our hybrid stochastic approach should integrate other available signals such as the dynamics and upregulation of homeostatic and inflammatory cytokines.

Several mechanistic assumptions and practical simplifications were made to approach the broader biological context of CAR T cell therapy. First, we assumed that immune reconstitution varies minimally across patients, but our results demonstrate that this approach contains sufficient feed-back to explain CAR spike and decay. A different approach would be to re-parameterize normal T cell dynamics over time, which could reveal whether variability in normal T cell growth is necessary to explain outcomes. Alternatively, CAR density over time could be driven by unknown external stimuli. These hidden processes could be used to explain the distribution of progression times, applying more complicated approaches that integrate time-dependence^44,45^.

Second, our model assumes that tumor cell proliferation is independent of tumor burden. However, the tumor growth rate might depend on tumor burden, compared to an innate carrying capacity. In this context, one could explore other sources of tumor burden variability that originate from a logistic dependence of tumor cell proliferation on tumor volume, called proliferation-saturation^46,47^. Future studies of the stochastic process should carefully consider non-linear relationships between tumor size and growth as a potential dynamic biomarker for the success of CAR T cell therapy.

Third, our probabilistic measure of PFS did not include the evolution of resistance to CAR T cell therapy by genetic or epigenetic mechanisms^9^. Our results can be seen as conditional on non-resistance, which would add an additional probabilistic modeling layer.

Fourth, detection of normal CD19+ B cells in circulation long after CAR T is likely evidence that functional CAR T cells no longer persist in the host. It is unclear whether B cells themselves are the driving event for continued persistence. Thus, we assume that non-tumor sources of CD19 do not play a role during the activity of CAR T cells.

Last, immune response against gene modified cytotoxic cells^48^ is a mechanisms we do not consider here. Cytotoxicity could result from rejection of the CAR expressing cells by anti-transgene response^49^, or anti-CAR T cell immune response against the murine single-chain variable fragment (scFv)^24^ that is incorporated in most CD19-targeting CARs. The complex nature of these immunogenicity-related rejection mechanisms renders it unlikely to generalize and model them at this point. In addition, immunogenic effects are unlikely following a single infusion with appropriate conditioning/lymphodepletion (i.e. fludarabine and cyclophosphamide)^50^, which is why we excluded them. Of note, these effects could play a more pronounced role in cases of multiple CAR T cell treatments.

With the mechanisms modeled here, cure occurred early, progression occurred late. This difference in timing of events indicates that rare patient events observed with cure way beyond 100 days are driven by other mechanisms, such as a vaccine effect. Mathematical dynamical models of such effects are lacking behind, pointing to promising new research avenues. Parts of our approach established here can serve a starting point to such efforts.

Complex relationships between tumor properties and second dose emerge because a single dose is already very efficient. Slow growing tumors will not improve much with a second dose. On the other hand, large tumors that would progress and are not treated twice within a short period of time will not show much improvement for a wide range of tumor growth rates. Hence, average and smaller tumors that grow at median or faster rates could see an improvement of 40% or higher, relative to the baseline single treatment.

CAR T cell therapy can cause severe cytokine release syndrome (CRS), or a characteristic immune cell-associated neurologic syndrome (ICANS). Both are associated with significant co-morbidity and mortality^51–53^, and are characterized by high levels of inflammatory cytokines and, to a lesser extent, by high numbers of CAR T cells in the blood^6^. Both forms of toxicity occur in predictable distributions across patient populations^54^. Published data from the ZUMA-1 trial demonstrates an association between CAR T cell expansion and grade 3-4 (severe) ICANS associated toxicities^6^. Elevation of key inflammatory cytokines (IFN-γ, IL-6, IL-1) are also associated with both severe ICANS and CRS. However, a homeostatic cytokine (IL-15) seen at higher levels in responding patients was also associated with toxicity. Several biomarkers were associated with severe ICANS associated toxicity, but not CRS (ferritin, IL2-Rα and GM-CSF)^55,56^. Thus, one could hypothesize that CAR T cell expansion and the occurrence of toxicities can be explained by modeling dynamic cytokine/biomarker levels in periphery together with CAR T cell repopulation dynamics and toxicity, in relation to homeostatic (IL-15) and inflammatory (IL-6, IFN-γ, IL-1) cytokines. Computational and mathematical tools, similar to the ones we developed here, could be used to calibrate such a model, assess parameter sensitivities, and make predictions of patient outcomes. Here, we present an important first step towards integration of these mechanisms into a quantitative understanding of CAR T cell therapy, modeling important complex mechanisms *in vivo*, and establishing that cellular immunotherapies can benefit from an ecological modeling perspective to establish mechanistic understanding of treatment failure and success.

## Acknowledgements

The authors would like to thank Andrew Stein (Novartis) and William Denney (Human Predictions LLC), as well as all members of the Altrock and Enderling labs at Moffitt Cancer Center for constructive comments and helpful discussions on an earlier version of this manuscript.

