## Supplementary Information and Materials for "Response to CAR T cell therapy can be explained by ecological cell dynamics and stochastic extinction events"

March 30, 2020

#### Contents

**1 Introduction** **2**

**2 Stochastic tumor cell extinction with a simple CAR-decay model** **2**

**3 Co-evolutionary dynamics among normal, CAR T, and tumor cells** **3**
3.1 Motivation . . . . . 4
3.2 Modeling re-emergence of normal T cells, CAR dynamics, and CAR-tumor interactions 5

**4 Data analysis** **8**

**5 Small fluctuations are relevant in the small tumor limit** **10**

### 1 Introduction

In this supplementary text, we lay out the theoretical foundations of or modeling approach, describe further details of the data analysis and model fitting procedures, and introduce and analyze a hybrid stochastic-deterministic nonlinear modeling approach used for efficient simulations and as a predictive tool.

First, we discuss a simple two dimensional stochastic extinction process that could be used to approximate tumor cell extinction as a result of a decaying CAR T cell population. However, this approach does not have the ability to describe the dynamics peak in CAR T cell density.

Second, we motivate and introduce a more comprehensive, four dimensional co-evolutionary framework in its mean-field limit. This framework can describe more complex features of both T and tumor cell populations during treatment, in particular CAR peak dynamics.

Third, we describe how we analyzed and integrated the available data using the four dimensional framework. The data used consists of (a) median T cell/lymphocyte counts in patients over time, (b) quartile CAR T cell densities in patient over time, and (c) estimates of median tumor burden (derived as tumor volume) over time, which were estimated based on the tumor burden of patients that had progressed at days 30, 60, 90 post CAR administration (beginning of treatment). The data analysis assumes fixed median initial cell densities, and we apply a nonlinear global optimization framework to minimize a pre-defined loss function. This minimization approach leads to possible model parameterizations and provides proof of principle that a four dimensional co-evolutionary framework can be used to describe CAR T cell therapy dynamics.

Last, we introduce a stochastic framework that corresponds to the mean-field model, and which can be used to describe stochastic tumor extinction events. These events are important, as the mean-field approach typically only shows transient tumor reduction–long-term tumor extinction is not stable in the mean-field. To effectively model the fully parameterized four dimensional dynamical system, we introduce and discuss a hybrid stochastic-deterministic framework.

### 2 Stochastic tumor cell extinction with a simple CAR-decay model

First, we discuss an approach that does not consider CAR T cell expansion or feedback by tumor antigen, but describes the essence of CAR T cell predation and provides an intuition for tumor cell extinction driven by CAR T cells. Let us consider two cell populations; CAR T effector cells $E(t)$ that kill tumor, and malignant B cell $B(t)$. In the simplest setting, we assume that tumor killing cells $E$ have their maximal value at time $t = 0$, and that these cells obey a simple death process. That is, they will go extinct with probability 1 eventually:

$$E \xrightarrow{d_E} \emptyset. \tag{1}$$

Meanwhile, these tumor killing cells contribute to tumor cell death, but the tumor follows a birth and death process

$$B \xrightarrow{b_B} B + B, \tag{2}$$

$$B \xrightarrow{d_B} \emptyset, \tag{3}$$

however, we would have to assume that the tumor cell birth rate is a function of $E(t)$. The simplest approach, however, would be to assume a time scale separation, and set $d_B = \delta_B \, E_0$. Then, we can write down the following master equation for the Markov process that governs tumor cell count over time (using $\dot{f} = df/dt$ notation)

$$\dot{P}_B(t) = b_B(B-1)P_{B-1}(t) + d_B(B+1)P_{B+1}(t) - (b_B + d_B)B\,P_B(t) \tag{4}$$

where $P_B(t)$ is the probability to find the system in state $B$ at time $t$. Using the generating function approach [1], we can obtain the following partial differential equation (PDE) for the generating function $F(t,x) := \sum_B P_m(t)\, x^B$ (whereby $\partial_x$ means partial derivative with respect to $x$)

$$\dot{F} = b_B(x^2 - x)\partial_x F + d_B(1 - x)\partial_x F \tag{5}$$

subject to boundary conditions $F(t,0) = P_{B=0}$, and $F(t,1) = 1$, and the initial condition $F(0,x) = x^{B_0}$. Here, $B_0$ is the initial tumor size, and $P_{B=0}$ is the tumor extinction probability, which turns out the be the quantity we are interested in. The PDE for $F$ is of Lagrange type and can be solved exactly

$$F(t,x) = \left( \frac{(x-1)d_B\, \mathrm{e}^{(b_B - d_B)t} - b_B\, x + d_B}{(x-1)b_B\, \mathrm{e}^{(b_B - d_B)t} - b_B\, x + d_B} \right)^{B_0}, \tag{6}$$

which then yields the probability of tumor extinction at time $t$:

$$P_{B=0}(t) = F(t,0) = \left( \frac{d_B - d_B\, \mathrm{e}^{(b_B - d_B)t}}{d_B - b_B\, \mathrm{e}^{(b_B - d_B)t}} \right)^{B_0}. \tag{7}$$

Since we made the assumption of a constant tumor death rate, we can now heuristically calculate the probability of cure as the conditional probability of two independent events, i.e. the product of $P_{B=0}$ and the probability that the CAR T cell population $E$ does *not* go extinct. For the simple death process assumed for $E$, we obtain the extinction probability

$$P_{E=0}(t) = \left(1 - \mathrm{e}^{-d_E\, t}\right)^{E_0}, \tag{8}$$

and thus could approximate the likelihood of tumor extinction, conditioned on CAR T cell survival, as $P_{B=0} \times (1 - P_{E=0})$.

However, in a more realistic setting, the tumor death rate is time dependent via its dependence on the number of tumor killing T cells, the CAR T cell birth rate is not zero, and its birth and death rates are also time dependent. This implies that one can find a higher-dimensional process that removes this explicit time-dependence [2]. In the following, we identify a four-dimensional system that models T and tumor cell co-evolution more comprehensively.

#### 3 Co-evolutionary dynamics among normal, CAR T, and tumor cells

A comprehensive mathematical model of tumor cell and CAR T cell evolution has to include non-monotonic changes of the total CAR T cell population, as well as the ability to adjust the relative contribution of CAR T cells that replicate, and CAR T cells that kill tumor at any point in time. This could be achieved by assuming that changes in the CAR T cell population are subject to a time-varying function. In the following, we develop a model that attributes such time dependence to a third T cell population, namely normal T memory cells.

To motivate a comprehensive approach modeling CAR T cell and tumor cell dynamics, we define the desired properties the model should contain. First, tumor growth should be countered by CAR T cell predation, such that if predation occurs at a higher rate, the tumor shrinks. Second, tumor, or B cell antigen (CD19) should affect CAR T cell expansion. Third, the CAR T cell product's volume (population size) and composition should be integrated, leading to a CAR T cell population that is sub-divided into at least two populations, for example (central) memory and effector CAR T cells. Fourth, the overall CAR T cell population should have the ability to expand and contract, both as a result of temporally changing signals within the patient.

##### 3.1 Motivation

We present here a game-theoretic, or co-evolutionary approach that leads to a mechanistic mathematical model, which can be used to describe the likelihood of successful treatment. As a key property, treatment success should be proportional to the CAR T cell population's ability to expand before its inevitable contraction. We contrast our approach to a simple statistical analysis that does not capture CAR expansion, see Figure 1 A, which shows the inner quartiles of the trial ZUMA-1 trial [3]. The characteristic maximum in observed CAR T cell concentration occurs about a week post-infusion. In addition, on can observe that three months post-infusion, the CAR population is still present in the peripheral blood in more than 50% of patients. This can be approached as a simple decay problem, and one can ask whether it is likely that exponential decay governs the dynamics, or whether other time scales matter. In Figure 1 B we compare the results (P-values) of linear regression analysis to the log-linear or log-log transformed median CAR T cell concentrations. In this context, it becomes apparent that exponential decay does not fully capture the dynamics. Rather, CAR T cells seem to exhibit a different half-life and long-term behavior; the long-term dynamics seem to be best approximated by a time dependence of the form $\sim t^{-\beta}$ (see Figure 1 C, D, E). This might indeed be indicative of some form of memory from the initial time point (or before), thus it highlights the role of the system's state before CAR administration.

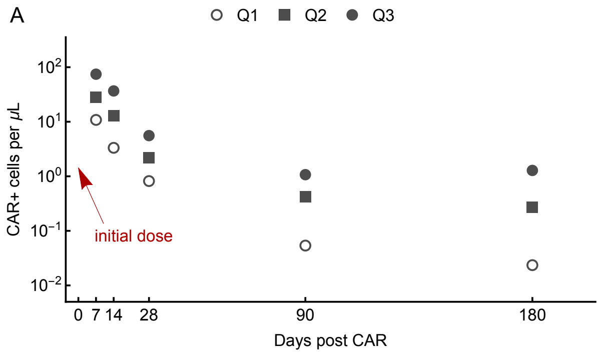

B

| | | | |
|---|---|---|---|
| Q1 | CAR($t$) | $\beta$ | P-value |
| | $\propto e^{-\beta t}$ | 0.0331 | 0.1530 |
| | $\propto t^{-\beta}$ | 1.9724 | 0.0004 |
| Q2 | CAR($t$) | $\beta$ | P-value |
| | $\propto e^{-\beta t}$ | 0.0245 | 0.0526 |
| | $\propto t^{-\beta}$ | 1.5185 | 0.0019 |
| Q3 | CAR($t$) | $\beta$ | P-value |
| | $\propto e^{-\beta t}$ | 0.0215 | 0.0940 |
| | $\propto t^{-\beta}$ | 1.3932 | 0.0089 |

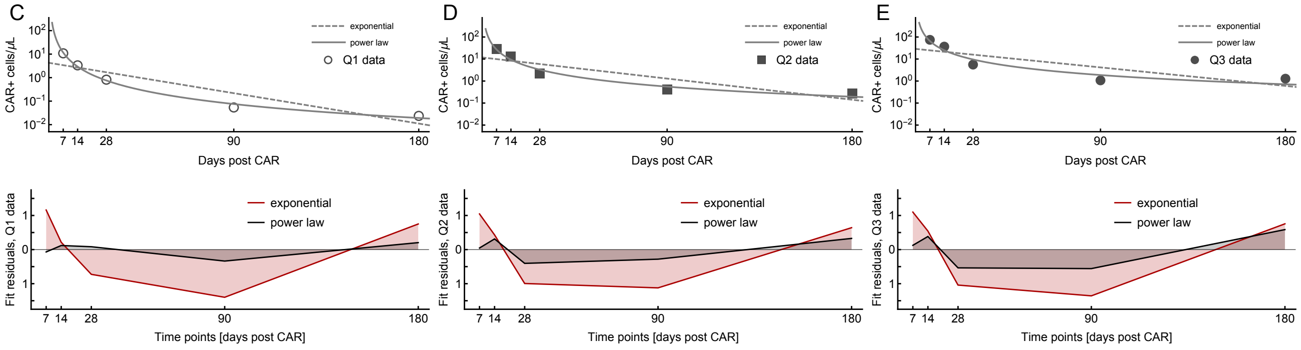

**Figure 1: A:** Quartile data of CAR T cell count in periphery reproduced from 101 patients of the ZUMA-1 trial [3] (lower quartile: circles, median: squares, upper quartile: discs). Of note, the initial median CAR T cell dose (at time 0) was 0.36 cell per $\mu$L blood–the fitting procedure presented here only accounts for the decay of CAR and can not describe its initial increase. **B:** We fit exponential and power law decay to the data (using *LinearModelFit* in Wolfram Mathematica), resulting in different values of the respective decay parameter $\beta$. Throughout, power law decay shows a significantly improved fit to these data points. **C:** Exponential ($\exp^{-\beta t}$, dashed) and power law ($t^{-\beta}$, solid) fits to the lower quartile data (circles) on the top, respective fit residuals on the bottom. **D:** Exponential ($\exp^{-\beta t}$, dashed) and power law ($t^{-\beta}$, solid) fits to the median data (squares) on the top, respective fit residuals on the bottom. **E:** Exponential ($\exp^{-\beta t}$, dashed) and power law ($t^{-\beta}$, solid) fits to the upper quartile data (disks) on the top, respective fit residuals on the bottom.

To motivate our assumptions, we first state the following observations:

- If lymphodepletion of normal T cells does not occur, the CAR T cells do not expand *in vivo*. **Implication:** *A carrying capacity, or other feedback of the overall lymphocyte count must exist, which is related to the total T cell count over time. Otherwise, CAR T cells would always expand.*
- The CAR T cells have an initial spike in concentration before decreasing and leveling out to a constant (which may or may not be zero). **Implication:** *The amount of space, or niche available to CAR T cells changes over time.*
- Influences of dynamic cytokine changes or other factors impact both normal and CAR T cells, and introduce a form of memory. **Implication:** *Some cellular reaction rates, such as the CAR T cells' growth rate, could be time dependent explicitly.* Note, however, that complete clinical response is possible with undetectable CAR T cell levels in the long term.

We here propose a mechanism that drives the early of CAR T spike, followed by decay. In this work we postulate that the mechanism that drives the dynamics of CAR cells is competition, or co-evolution, with normal T cells, as well as the dynamics feedback of the targeted tumor cell population.

#### 3.2 Modeling re-emergence of normal T cells, CAR dynamics, and CAR-tumor interactions

Our approach is motivated by two key observations. First, chemo-depletion of T cells seems to be required for proliferation of CAR T cells. This observation implies that there is an innate carrying capacity shared by both types of T cells (wild-type and CAR). The initial depletion is necessary to give CAR cells an ability to proliferate. The nonlinear nature of this dynamic behavior indicated that there are compartments of CAR T cells that play distinct roles, such as naïve, central memory and effector cells.

Second, after a peak at around 7 days post injection, CAR levels begin to decay. This observation could imply that a wild-type/normal T cell population has an overall advantage–otherwise the long-term levels of CAR would not tend to zero. These observations lead to the following model assumptions.

##### Assumptions

We conceptualized a model of CAR T cell therapy based on four key assumptions. First, we consider two sub-populations of CAR T cells. We combine the compartments of naïve and central memory cells into one, which we call "memory" CAR T cells, $M$. When presented with antigen, $M$ cells differentiate into a second compartment of effector and effector memory CAR T cells, which we call "effector" CAR T cells, $E$. These cells do not divide, and target and kill antigen presenting tumor cells, $B$.

Second, we introduce co-evolution, or competition, in the naïve/memory T cell compartment. To this end we consider the normal T cell population, $N$. The populations $M$ and $N$ can inhibit each others' expansions, but at different rates. That is, we assume negative feedback mechanisms between $M$ and $N$, their net growth rates depend on the total amount $M + N$, for which one can introduce carrying capacities ($K_M < K_N$). In this sense, CAR T cells are maladapted: normal T cells have the propensity to expand to higher numbers during immune reconstitution. Hence, memory CAR T cells eventually will be selected against (unless other long-term memory mechanisms would be considered) and decay, but may get a head start and expand, due to lymphodepletion. This expansion is modeled by asymmetric differentiation of $M$ cells into $E$ cells.

Third, we assume that asymmetric differentiation from memory to effector CAR T cells is antigen-dependent. CAR effector cells kill tumor cells at a high rate, and at the same time these effector cells have a finite life-span. We introduce a predator-prey interaction term between $E$ and $B$ that defines the killing rate of the tumor cells when engaged by a CAR effector cell. For simplicity, we assume a linear (Holling type I) predation rate. Other, more complex interaction terms could be considered at the cost of additional parameters.

Last, we assume simple exponential growth of the tumor cell population $B \propto e^{r_B t}$. These assumptions and relationships can be cast as the following system ordinary differential equations (ODEs):

$$\dot{N} = N(r_N - a_{11}N - a_{12}M), \tag{9a}$$
$$\dot{M} = M(r_M - a_{21}N - a_{22}M), \tag{9b}$$
$$\dot{E} = r_E(B)M - \gamma_E BE - d_E E, \tag{9c}$$
$$\dot{B} = r_B B - \gamma_B BE. \tag{9d}$$

With initial values of $N_0,\ M_0,\ E_0,\ B_0$, respectively. Here $r_M$ and $r_N$ are the net growth rates of CAR and normal memory T cells, respectively. The coefficients $a_{ij}$ are the interaction rates that determine the magnitude of negative feedback that the $j$ subtype has on the $i$ subtype. $r_E$ is the asymmetric differentiation rate and $d_E$ is the CAR effector death rate. The rates $\gamma_{E,B}$ are the effector-tumor interaction rates, where an effector or tumor cell dies upon interaction, respectively. Finally, $r_B$ is the tumor's intrinsic net growth rate.

##### Carrying capacity formulation

A general formulation of interaction involves a competitive Lotka-Volterra model with interaction coefficients $a_{ij}$. However, it is very difficult to infer these parameters from biological data. A simplifying assumption involves scaling these coefficients $a_{ij} = 1/K_i$, i.e. introducing a carrying capacity that each subtype would have in the absence of the other subtype. Hence, by redefining the parameters $a_{ij} = r_i/K_i$, we arrive at

$$\dot{N} = r_N N \left[1 - \left(\frac{N+M}{K_N}\right)^{\beta}\right], \tag{10a}$$
$$\dot{M} = r_M M \left[1 - \left(\frac{N+M}{K_M}\right)^{\beta}\right], \tag{10b}$$
$$\dot{E} = r_E(B)M - \gamma_E BE - d_E E, \tag{10c}$$
$$\dot{B} = r_B B - \gamma_B BE. \tag{10d}$$

Preliminary optimization of the model parameters showed that $\beta \to 0$, and that $r_i$ is large. Rescaling $r_i = r_i/\beta$ $(i = N, M)$ and letting $\beta \to 0$ we obtain the following form for our model

$$\dot{N} = -r_N N \ln\left(\frac{N+M}{K_N}\right), \tag{11a}$$
$$\dot{M} = -r_M M \ln\left(\frac{N+M}{K_M}\right), \tag{11b}$$
$$\dot{E} = r_E(B)M - \gamma_E BE - d_E E, \tag{11c}$$
$$\dot{B} = r_B B - \gamma_B BE, \tag{11d}$$

which is the form of the mean-field model we use for the remainder of the text.

**Steady states**

Since $N$ and $M$ are only dependent on each other, we can consider them separately. If $M^* = 0$, then $N^* = 0, K_N$. Suppose $M^* \neq 0$, then if $N^* = 0$ we have $M^* = K_M$. A coexistence state is only observed in the case $K_N = K_M$. Since indefinite persistence of CAR T cells was not observed clinically, we conclude that $K_N > K_M$ is clinically/biologically realistic. This assumption helps to limit the parameter region.

We note that there are *no* coexistence states. The stability and states can be summarized as follows:

- **Unphysical solution**: $(0, 0, 0, e^{r_B t})$ - the T cell-absent state. This state is *always* unstable, but unphysical in the sense that there would have to be an absence of normal T cells at any time.

- **Failed treatment**: $\left(K_N, 0, 0, e^{r_B t}\right)$ - the CAR cells are depleted and the tumor is not eliminated, leading to exponential tumor growth. This state is stable if $K_N > K_M$.

- **Cured patient**: $\left(0, K_M, \frac{K_M r_E(0)}{d_E}, 0\right)$ - the CAR cells eliminate the tumor and outcompete the wild-type to become the resident population. This case os also unrealistic–we expect the normal T cells to have an innate advantage due to a deeper stem and progenitor cell pool of normal cells. This state is stable if
$$K_M > \max\left(K_N, \frac{d_E r_B}{\gamma_B r_E(0)}\right).$$

- **Stable patient**: $\left(0, K_M, \frac{r_B}{\gamma_B}, B^*_{\text{stable}}\right)$, where $B^*_{\text{stable}}$ satisfies
$$r_E(B^*_{\text{stable}})\gamma_B K_M - r_B(\gamma_E B^*_{\text{stable}} E + d_E) = 0. \tag{12}$$
Here, the CAR-tumor interactions dynamically stabilize the tumor. This case is also unrealistic–we expect the wild-type population to be present. Nonetheless, this state is stable if
$$K_M > \max\left(K_N, \frac{\gamma_E r_B}{\gamma_B r'_E(B^*_{\text{stable}})}\right).$$

- **Tumor growth**: This state is slightly different than the the states considered so far in that it is not a "fixed point". Ultimately, we are concerned with tumor growth. If the maximum value of $E$ is such that $\dot{B} > 0$, then we have an unsuccessful treatment outcome. Hence, if $E(t) < E_{\max} = \frac{r_B}{\gamma_B}$ the tumor will grow (note that this bounds solutions away from the stable patient outcome).

We can see that under the assumption $K_N > K_M$, the only long-term states are failed treatment and eventual tumor growth. Note that this includes cases in which the tumor shrinks temporarily. In the deterministic setting considered so far, the tumor cell population can become arbitrarily small and spend long times in a regime near 0, where random cell death events could lead to the elimination of tumor. These small-population size effects cannot be adequately captured with a mean-field ODE model. Hence, we will eventually turn to a stochastic model formulation. Before we discuss the stochastic system equivalent to the mean-field dynamical system, we first ask under which conditions CAR T cell densities can exhibit non-monotonic temporal behavior.

##### Conditions for non-monotonic CAR T cell population dynamics

In Figure 1 A, we see that the CAR levels reach a maximum if we factor in the initial CAR density, followed by decay. We here derive the conditions needed for this peak to occur, in the general case, and then for the carrying capacity model as a corollary.

Let us first consider the CAR memory compartment. For a spike in CAR to occur, we require there to exist an $N_{\text{peak}}, M_{\text{peak}}$ such that $\dot{M} = 0$ and $\ddot{M} < 0$. This leads to

$$M_{\text{peak}} = \frac{r_M - a_{21} N_{\text{peak}}}{a_{22}}. \tag{13}$$

Differentiation of Eq. (9b) leads to

$$\ddot{M} = \dot{M}(r_M - a_{21}N - a_{22}M) - M(a_{21}\dot{N} + a_{22}\dot{M}). \tag{14}$$

Evaluating this at $N_{\text{peak}}, M_{\text{peak}}$ leads to

$$\ddot{M} = -a_{21}\dot{N}M_{\text{peak}}. \tag{15}$$

Hence, a peak will exist if $\dot{N} > 0$ at that point. Furthermore, we require $M_{\text{peak}} > 0$, which implies $r_M > a_{21}N_{\text{peak}}$. Looking at Eq. (9a) at the peak, yields

$$\dot{N} = N\left(r_N - \frac{N_{\text{peak}}\Delta + a_{12}r_M}{a_{22}}\right), \tag{16}$$

where $\Delta = a_{11}a_{22} - a_{21}a_{12}$ is the determinant. This leads to the requirement that

$$a_{22}r_N > N_{\text{peak}}\Delta + a_{12}r_M. \tag{17}$$

In the carrying capacity case, we note that $\Delta = 0$ and so our conditions reduce to $K_N > K_M > N_{\text{peak}}$. We recognize the first condition as the requirement that the CAR memory only state is unstable.

#### 4 Data analysis

We used the median, lower and upper and quartiles of 101 patients from the ZUMA-1 trial, as published by Neelapu *et al.* (2017) [3]. This data set contains total peripheral CAR T cell concentrations (quantiles), from peripheral blood measurements, recorded at days 7, 14, 28, 90, and 180 post CAR injections. These 15 data points, in combination with normal T cell/lymphocyte counts obtained from some of these patients independently at Moffitt Cancer Center at day 0, 5, 7, 14, 28, 90, 180, as well as using clinical response data as a proxy for tumor size, were used to fit the parameters of the T cell-CAR T cell and tumor cell dynamics model in its deterministic limit.

To arrive at tumor size estimates, we assumed the following breakdown of clinical responses, respectively at time $t = 30$, 60 and 90 days post CAR injection. Complete response (CR) was counted as tumor size $B(t) = 0$/no detectable tumor. Stable disease (SD) was counted as the initial tumor size, $B(t) = B_0$. Progressive disease (PD) was counted as twice the initial tumor size, $B(t) = 2 \times B_0$.

Note also that the breakdown of the total CAR T cell population into compartments $M$ and $E$ was only available at time of CAR administration. Hence, when fitting these data, one must either make assumptions on the phenotype breakdown at each available time point, or compare the model solution $(M + E)(t)$ concentration against the total CAR T data. We separated the data into subsets assuming fixed fraction of CAR memory cells $f_{\text{mem}} = M/(M + E)$, and thus were able to treat the compartments $M$ and $E$ as separate data/modeling functions.

To fit the data, we implemented a BFGS constrained optimization routine using Julia's built in optimization and differential equation solvers to determine the optimal parameters which minimize our predefined loss function [4, 5]. We used a weighted-least squares function, where the weights are the variance of the data at each time point. Using the quartile data only, we assume that the distribution of CAR T cell levels at each time point is normally distributed about the median and calculate an estimate of the variance from the interquartile range:

$$\sigma^2 \approx \left(\frac{Q_3 - Q_1}{1.35}\right)^2 . \tag{18}$$

We first fit the normal and memory CAR T cells, since the two compartments $M$ and $E$ are decoupled from the other two ($E$, which depends on $M$ and $B$, and $B$, which depends on $E$). The loss function used is

$$\theta_{N,M} = \lambda_0 \sum \frac{[M(t_i) - \hat{M}_i]^2}{\sigma_{M,i}^2} + \lambda_1 \sum [N(t_i) - \hat{N}_i]^2, \tag{19}$$

where the $\lambda$'s are tunable weights that can be used to adjust the loss function landscape. Different $\lambda$'s lead to different optimal parameter sets. We generated trajectories by using the three quartiles and using the assumption that the memory fraction $f_{\text{mem}}$ was fixed over time. We chose $f_{\text{mem}} = [0.1, 0.5, 0.9]$, which led to a total of nine trajectories.

Of note, the initial CAR T cell spike near day 7, followed by a fast deacy, can be well captured by three fitted parameters $r_N, r_M, K_M$, in principle. However, the data showed a slowing-down of CAR decay after day 14–some patients still had observable CAR levels 6 months post-infusion. This behavior could not be captured with these three parameters only. To account for this, we introduced two additional parameters that govern a switch in the value of the CAR growth rate $r_M$. As $K_M$ is intricately linked to the long-term stability of $M$ and $N$, it made more sense to put this time-dependence into $r_M$. We introduced the function

$$r_M(t) = \frac{r_{M,\text{max}} - r_{M,\text{min}}}{1 + e^{t-\tau}} + r_{M,\text{min}}. \tag{20}$$

Many functional forms can be given for the antigen/tumor size-dependent production rate of effector CAR T cells, $r_E(B)$. Clinical trial data and *in vitro* experiments show that more antigen presentation leads to an increase in differentiation into effector cells [6, 7, 8]. We modeled this as the following piecewise-linear function

$$r_E(B) = r_E(0) \left[1 + \alpha_1 \min\left(\frac{B}{B_0}, \alpha_2\right)\right]. \tag{21}$$

To obtain the parameters in the $E$ and $B$ compartment, we introduced a second loss function

$$\theta_{E,B} = \lambda_0 \sum \frac{[E(t_i) - \hat{E}_i]^2}{\sigma_{E,i}^2}, \tag{22}$$

where the fitted parameters for $M, N$ are used to solve for the remaining parameters.

| Biological parameter | Symbol | Fitted model | Stochastic simulation model | Reference |
|---|---|---|---|---|
| Normal T cell carrying capacity | $K_N$ | $5.00 \times 10^2$ cells/$\mu$L | $2.50 \times 10^{11}$ cells | Turtle *et al.* (2016) [9] |
| CAR memory T cell carrying capacity | $K_M$ | $1.60 \times 10^2$ cells/$\mu$L | $8.00 \times 10^{10}$ cells | this work |
| Normal net growth rate | $r_N$ | $1.60 \times 10^{-1}$ day$^{-1}$ | unchanged | this work |
| Initial CAR memory net growth rate | $r_{M,\max}$ | $4.14 \times 10^{-1}$ day$^{-1}$ | unchanged | this work |
| Final CAR memory net growth rate | $r_{M,\min}$ | $2.15 \times 10^{-2}$ day$^{-1}$ | unchanged | this work |
| CAR memory net growth rate switch | $\tau$ | $1.88 \times 10^1$ days | unchanged | this work |
| Basal CAR asymm. diff. rate | $r_E(0)$ | $2.26 \times 10^0$ day$^{-1}$ | unchanged | this work |
| Effector death rate | $d_E$ | $2.93 \times 10^{-1}$ day$^{-1}$ | unchanged | this work |
| Tumor growth rate | $r_B$ | $6.50 \times 10^{-2}$ day$^{-1}$ | unchanged | Roesch *et al.* (2014) [10] |
| Effector exhaustion rate | $\gamma_E$ | $3.00 \times 10^{-3}$ days$^{-1}$*cm$^{-3}$ | $3.00 \times 10^{-12}$ (days*cells)$^{-1}$ | this work |
| Tumor killing rate | $\gamma_B$ | $3.00 \times 10^{-2}$ (days*cells/$\mu$L)$^{-1}$ | $6.00 \times 10^{-11}$ (days*cells)$^{-1}$ | this work |
| Increased asymm, diff. rate | $\alpha_1$ | $4.29 \times 10^0$ | unchanged | this work |
| Max tumor ratio | $\alpha_2$ | $1.35 \times 10^0$ | unchanged | this work |
| Initial ALC | $N(t=0)$ | 6.00 cells/$\mu$L | $3.00 \times 10^9$ cells | Locke *et al.* (2017) |
| Initial CAR | $M(0)+E(0)$ | 0.36 cells/$\mu$L | $1.80 \times 10^9$ cells | Locke *et al.* (2017) |
| Initial tumor | $B(0)$ | 200 cm$^3$ | $2.00 \times 10^{11}$ cells | Locke *et al.* (2017) |

**Table 1:** Median parameter values of our model, using minimization of the loss functions (19) and (22). Model fitting used T cell densities in cells per $\mu$L (peripheral) and tumor size in cubic cm. Stochastic hybrid simulation modeling used transformation to cell counts, assuming that every patient has on average 5L of blood that contains 1% of the T or CAR T cell population, and that 1 cm$^3$ contains $10^9$ tumor cells on average. Initial conditions are the median values. ALC: absolute lymphocyte count.

In summary, the mean-field ODE model was used for this non-linear constrained optimization approach to find best fits (of which there are potentially many). The parameter values obtained are shown in Table 1. In the next section, we introduce a stochastic framework that considers individual cellular events. We used the model parametrization presented in Table 1, for which some of the values had to be scaled to represent individual cellular events, as indicated by the additional column.

### 5 Small fluctuations are relevant in the small tumor limit

We have discussed the stability and fixed points of Eq. (9). This system describes initially successful treatment and a peak in CAR concentration over time. However, as normal lymphocytes reemerge and reaches a homeostatic level, CAR decays, the tumor eventually grows back and relapse occurs deterministically in the mean-field framework. Although this scenario is plausible, it is still one of several outcomes.

Complete durable responses have been observed in the clinic (typically without detectable CAR levels long-term). When the tumor population becomes small, stochastic effects (fluctuations) become relevant–the tumor might be subject to an extinction vortex [11]. These chance extinction events can lead to tumor extinction. Of note, our modeling framework implies that stochastic extinction is a *necessary* requirement for durable response, if CAR T cells do not persist indefinitely. In what follows, we will propose a stochastic framework that in the large population limit will converge to the deterministic dynamics.

##### 5.1 Stochastic dynamics

Let $N$, $M$, $E$, $B$ now be the cell numbers of wild-type, CAR memory, CAR effector and tumor cell populations, respectively. We define the following birth events,

$$N \xrightarrow{r_N} N + N, \tag{23}$$

$$M \xrightarrow{r_M} M + M, \tag{24}$$

$$M \xrightarrow{r_E(B)} M + E, \tag{25}$$

$$B \xrightarrow{r_B} B + B, \tag{26}$$

and the following death events

$$N \xrightarrow{\delta_N(N,M)} \emptyset, \tag{27}$$

$$M \xrightarrow{\delta_M(N,M)} \emptyset, \tag{28}$$

$$E \xrightarrow{\delta_E(B,E)} \emptyset, \tag{29}$$

$$B \xrightarrow{\delta_B(B,E)} \emptyset. \tag{30}$$

We can now calculate the transition rates to move state $i$ to state $j$. We choose a time interval small enough such that only a single event occurs (all other events occur on the order $O(\Delta t)^2$). Defining $T(N \pm 1, M, E, B | N, M, E, B) = T_N^{\pm}$ and noting that $r_i$ can depend on other populations we have the population transition rates

$$T_i^+ = r_i i, \quad i = N, M, E, B, \tag{31}$$

$$T_N^- = \delta_N N = r_N N \ln\left(\frac{N+M}{K_N}\right), \tag{32}$$

$$T_M^- = \delta_M M = r_M M \ln\left(\frac{N+M}{K_M}\right), \tag{33}$$

$$T_E^- = \delta_E E = \gamma_E E B + d_E E \tag{34}$$

$$T_B^- = \delta_B B = \gamma_B E B. \tag{35}$$

Denoting $P(N-1, M, E, B, t) = P_{N-1}$, the corresponding master equation is given by

$$\frac{\partial P}{\partial t} = T_{i-1}^+ P_{i-1} + T_{i+1}^- P_{i+1} - (T_i^+ + T_i^-)P, \tag{36}$$

where $i = N, M, E, B$.

A shortcoming of an exact simulation of this four dimensional stochastic framework is the computing time required for typical tumor size and lymphocyte count, which are in the orders of millions to billions. The number of events to observe a complete response is when the number of tumor cells approaches 0. The minimum number of stochastic individual cellular events needed then is clearly $B_0$, where $B_0$ is the initial number of tumor cells (see Table 1).

In practice, the total number of simulation events needed to approach tumor extinction will exceed $B_0$ by a substantial amount. On conventional CPUs, the amount of time it takes for a single run can vary from hours to days. The issue is that the transition rates are so high, which means that the time to a possible next event are very short. A way to overcome this obstacle was proposed by Gillespie himself, who suggested the method known as "tau-leaping" [12]. This method is similar to a forward Euler method for a continuous system, except that the update is taken as a Poisson distributed random variable. The time $\tau$ is given as a fixed step-size, which alleviates the events occurring rapidly, but error is introduced as the simulation is no longer exact (populations are assumed to be constant in the time interval $[t, t+\tau]$. This method works well in many cases, unless if the cell populations exhibit substantially different scales.

Suppose there are two populations, one small (stochastic fluctuations are relevant) and the other very large (fluctuations are not relevant). In this case, tau-leaping could potentially lead to large error if a step is too large, but could be computationally costly if it is too small. The issue is that one population's update rates occur on vastly different scales than the other's, hence the timing of respective events occur on separate scales. This issue is analogous to the problem where our tumor population decreases rapidly, but both the normal and CAR T cell populations remain relatively high for some time. Therefore, we here propose a method that exploits this separation of time scales, to produce a fast algorithm, while maintaining the fluctuations necessary for complete response.

##### 5.2 Hybrid model

Hybrid models that combine the speed of deterministic models with the accuracy of stochastic models has been used before in several different contexts [13, 14, 15]. Here, we develop a hybrid model for our four population system. The process involves communicating information about the population between both models and using them when appropriate. We will need to keep track of the discrete and continuous population sizes (to convert from continuous to discrete, we use the ceiling function). The final piece needed to connect the two is to define the threshold at which one considers fluctuations important. We set the stochastic threshold to be 100 cells. An outline of our procedure is given below:

- Define patient-level parameters (e.g. tumor growth rate, initial levels of CAR and normal T cells)
- Initialize the discrete and continuous population vectors.
- Set $t = 0.0$ and define the final time to run the simulation (we typically used $t = 300 - 1000$ measured in days).
- While $t < t_{\text{final}}$:
- Solve the ODE forward in time until the tumor population goes below the stochastic threshold (e.g. 100 cells) or the final time is reached.
- Update the new continuous and discrete populations.
- If a discrete (stochastic) event occurred, update the populations.
- Determine the time till next *stochastic* event $\tau$ and simulate the deterministic system until this next event (or $t_{\text{final}}$) is reached.

This method allows us to utilize the speed of simulating an ODE for the large populations and allows us to catch the complete response observed when the tumor population stochastically goes extinct.

##### 5.3 Parameter variability, virtual patient cohorts, and stochastic threshold sensitivity

Using the parameter values reported in Table 1, we created additional parameter variability in the form of a mixture model/perturbation parameter $\sigma$: We assumed that each parameter varies for each patient around the median with a normal distribution with mean equal to the median and variance equal to $\sigma$ times median.

To generate survival data in Figures 2, 3, and 4 of the main text, we ran the hybrid model for 1000-10000 computer-simulated instances, each with a potentially different set of parameters chosen from a distribution around the median parameter value. We then recorded the time and rate of cure or progression for each patient and parameter set. The Kaplan-Meier (K-M) curves were generated in *Wolfram Mathematica*, using the function *SurvivalModelFit*.

The perturbation parameter $\sigma$ that governed the variability of all fitted parameters. In another sense, $\sigma$ can be regarded as a sensitivity analysis parameter. Consider a parameter $\alpha$, then the perturbed values can be in the range $\alpha[1 + \sigma U(-1, 1)]$, where $U(a, b)$ is a uniform distribution with bounds $a, b$. From a clinical trial standpoint, we can consider $\sigma$ as a crude measure of patient variability, however without individual patient resolution. For example, if we wish to simulate a *single* patient to determine that individual's 'optimal' treatment as predicted from the median set of parameters fitted, we would choose $\sigma = 0$. If we want to generate a *cohort*, we consider $\sigma > 0$. For most analyses, we chose a range of $\sigma = 0.05 - 0.15$.

The hybrid model adds an additional exogenous parameter to the system–the stochastic threshold $S$. Clearly, as $S \to 0$, the system approaches the full ODE model, while if $S \to \infty$ the system will be fully stochastic. In our simulations we chose $S = 100$, but we investigated the sensitivity of the results for different $S \in [10, 1000]$ and found that our results are robust to the threshold, making the hybrid model a valid approach for solving these problems without compromising speed by using a fully stochastic model. To test this, we generated a virtual cohort of 1000 patients and compared the rate of cure, mean time of cure and progression distributions against different values of $S$.

We also calculated the mean cure time and progression time as a function of $S$. From this we obtained the coefficient of variations for cure $c_{\text{cure}} = \sigma/\mu \sim 0.036$ and $c_{\text{progression}} \sim 0.018$. This shows that the variability due to changing $S$ changed these quantities by 1.8 - 3.6% of their respective means.

The empirical distributions were also tested using the non-parametric K-S and Pearson tests. We considered $S = 100$ our null distribution and compared it against all the other distributions (both cure and progression time). All comparisons and both tests yielded p-values above 5%, with a median p-value of around 50%. This is a strong indicator that these came from the same underlying distribution. Thus we can reasonably conclude that the results are not sensitive to the choice of $S$.
